# Dysregulated *SASS6* expression promotes increased ciliogenesis and cell invasion phenotypes

**DOI:** 10.1101/2024.01.29.576599

**Authors:** Eleanor Hargreaves, Andrew D Jenks, Adina Staszewski, Athanasios Tsalikis, Raquel Bodoque, Mar Arias-Garcia, Yasmin Abdi, Abdulaziz Al-Malki, Yinyin Yuan, Rachael Natrajan, Syed Haider, Thomas Iskratsch, Won-Jing Wang, Susana Godinho, Nicolaos J Palaskas, Fernando Calvo, Tobias Zech, Barbara Tanos

## Abstract

Centriole and/or cilia defects are characteristic of cancer cells and have been linked to cancer cell invasion. However, the mechanistic basis of these effects is unknown. Spindle assembly abnormal protein 6 homolog (SAS-6) is essential for centriole biogenesis and cilia formation. In cycling cells, SAS-6 undergoes APC^Cdh1^-mediated targeted degradation by the 26S proteasome at the end of mitosis. Little is known about the function of SAS-6 outside of centrosome biogenesis. To examine this, we expressed a non-degradable SAS-6 mutant (SAS-6ND). Expression of SAS-6ND led to an increase in ciliation and cilia-dependent cell invasion, and caused an upregulation of the YAP/TAZ pathway. YAP/TAZ or ciliogenesis inhibition prevented SAS-6-induced invasion. SAS-6ND caused increased actin alignment and stress fiber coherency, and nuclear flattening known to promote YAP nuclear import. Finally, data from The Cancer Genome Atlas showed that SAS-6 overexpression is associated with poor prognosis in various cancers. Our data provide evidence for a defined role of SAS-6 in cancer cell invasion and offers mechanistic insight into the role of YAP/TAZ in this cilia-sensitive process.

**Synopsis:** 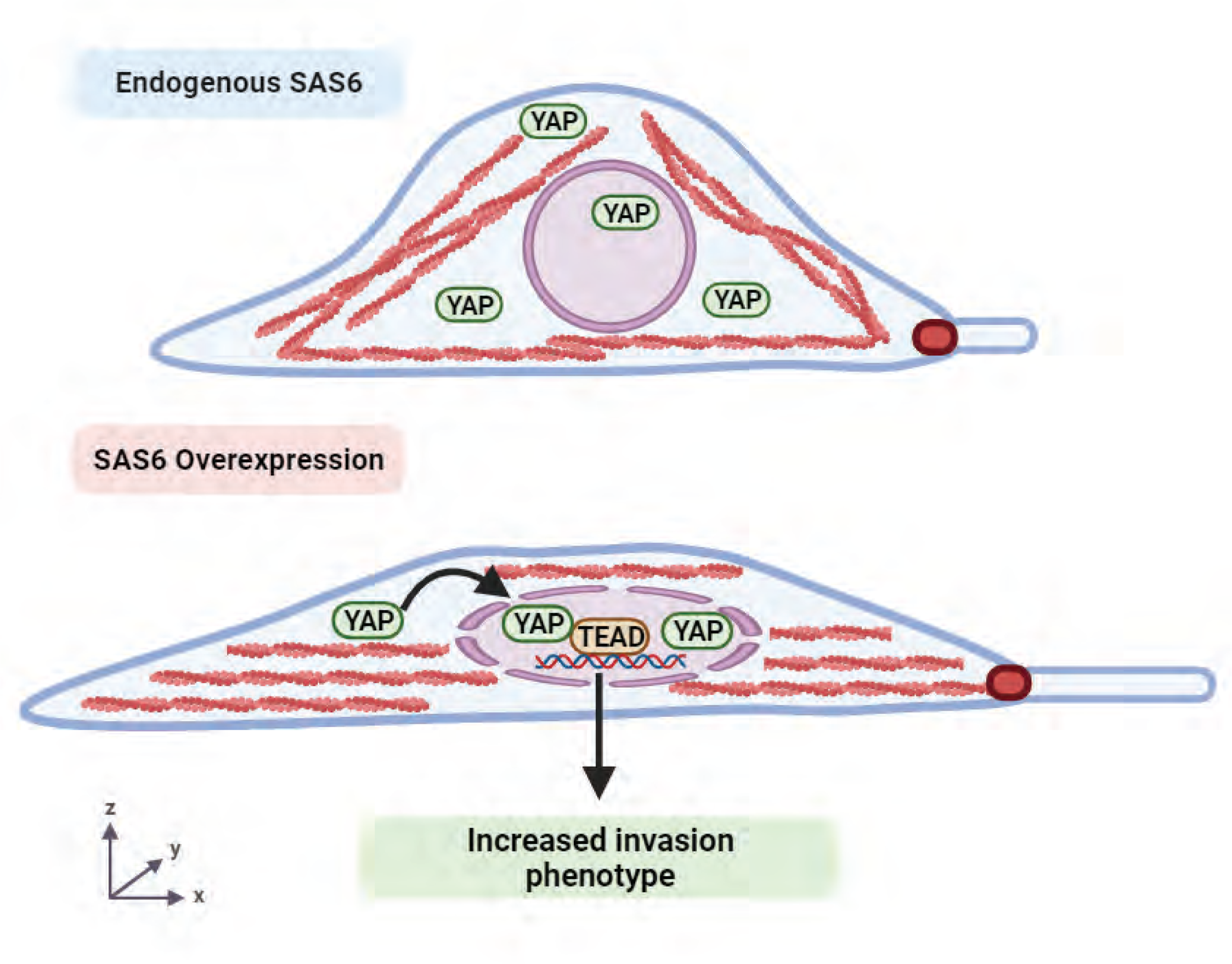

SAS-6 overexpressing cells show increased ciliation, actin cytoskeleton reorganization, cell flattening, YAP pathway activation and increased invasion

## Introduction

Centrosome and cilia abnormalities have been shown to be involved in cancer progression (Godinho & Pellman, 2014; Godinho *et al*, 2014; Han *et al*, 2009; Jenks *et al*, 2018; Wong *et al*, 2009), and in breast cancer cell invasion(Godinho *et al*., 2014).

Spindle assembly abnormal protein 6 homolog (*SASS6* gene, SAS-6 protein), is a key factor in centriole assembly and duplication (Leidel *et al*, 2005), a process that involves a sequence of coordinated events involving a set of six proteins (Delattre *et al*, 2006; Pelletier *et al*, 2006; Sugioka *et al*, 2017). SAS-6 together with SAS-5/STIL form the core of the cartwheel structure, a geometric scaffold that defines procentriole radial symmetry (Kitagawa *et al*, 2011) (Nakazawa *et al*, 2007) (van Breugel *et al*, 2011). The presence of the cartwheel is essential for the 9-fold symmetry organization of centrioles, as basal bodies lacking this structure are fragmented and disorganized (Nakazawa *et al*., 2007). The SAS-6 structure displays an N-terminal head domain, a coiled-coil domain that assists with dimerization and a C-terminal domain (van Breugel *et al*., 2011). SAS-6 dimers oligomerize via the N-terminal heads to form the nine-fold cartwheel which will also anchor microtubules (Kitagawa *et al*., 2011; van Breugel *et al*., 2011) as they elongate unidirectionally and form centrioles (Leidel *et al*., 2005; van Breugel *et al*., 2011).

Although the function of SAS-6 in the context of centrioles has been well studied, whether SAS-6 has any additional biological functions has not been explored. Polo-like kinase 4 (PLK4), a kinase that phosphorylates STIL to promote STIL and SAS-6 recruitment to the cartwheel structure (Dzhindzhev *et al*, 2014; Moyer *et al*, 2015; Ohta *et al*, 2014) (Arquint & Nigg, 2016), is deregulated in cancer (Godinho *et al*., 2014). PLK4 leads to centrosome amplification, resulting in the formation of invasive structures in breast cancer models (Godinho *et al*., 2014) as well as increased small vesicle secretion (Adams *et al*, 2021). Work by Shinmura et al showed that *SASS6* overexpression was associated with mitotic chromosomal abnormalities and poor prognosis in patients with colorectal cancer (Shinmura *et al*, 2015). *SASS6* is overexpressed in a number of human cancers, including kidney cancer, bladder cancer and breast invasive carcinoma (Shinmura *et al*., 2015) and reportedly promotes proliferation by inhibiting the p53 signaling pathway in esophageal squamous carcinoma cells (Xu *et al*, 2020). Interestingly, knockdown of *SASS6* reduced the growth of MDA-MB-231 triple-negative breast cancer cells (Du *et al*, 2021).

Studies of the function of SAS-6 in cancer are complicated because SAS-6 is periodically degraded at the end of mitosis/G1 by the APC-Cdh1 complex via its KEN box (Strnad *et al*, 2007). Here, we overexpressed a non-degradable SAS-6 KEN box mutant (SAS-6-ND) which is expressed throughout the cell cycle. SAS-6 overexpression led to an increase in cilia numbers and cilia length. Analysis of The Cancer Genome Atlas (TCGA) showed that SAS-6 overexpression was consistent with poor prognosis in adrenocortical carcinoma, low grade glioma, kidney, liver and lung cancer patients. This suggested a potential role for SAS-6 in metastatic cancer. Consistently, SAS-6 overexpression showed increased invasion that could be suppressed by blocking ciliogenesis. Transcriptome analysis revealed an upregulation of the YAP/TAZ pathway following expression of SAS-6ND. Notably, blocking YAP/TAZ function reverted *SASS6*-induced invasion. Analysis of cell morphology in SAS-6 overexpressing cells showed cell flattening and nuclear deformation, which causes opening of the nuclear pore complex (NPC) and YAP nuclear import (Elosegui-Artola *et al*, 2017). Our work shows a unique novel function of SAS-6 in invasion through the regulation of YAP/TAZ and provides rationale for interrogating the therapeutic potential of targeting *SASS6* as a potential strategy to prevent metastatic disease.

## Results

### SAS-6 promotes an increase in ciliogenesis

SAS-6 is an integral component of centrioles, and it is a key protein in centriole duplication. In cycling cells, SAS-6 is degraded in G1 by the APC^Cdh1^ complex (Strnad *et al*., 2007). However, a mutation in the conserved KEN box domain (Figure 1A), results in the stabilization of the protein (SAS-6 non degradable – ND-). It was previously shown by the Tsou laboratory that a SAS-6 mutant lacking the KEN box can localize to the mother centriole in G1 (Fong *et al*, 2014). Furthermore, in terminally differentiated cells of respiratory epithelia and in unicellular eukaryotes, SAS-6 localizes to basal bodies (Kilburn *et al*, 2007; Vladar & Stearns, 2007). Thus, we sought to understand whether SAS-6 would have any additional functions besides centrosome duplication in S-phase. To do this, we stably transduced RPE-1 cells with a SAS-6ND mutant under the regulation of a tetracycline-inducible promoter (Fong *et al*., 2014) (Figure 1B). As expected, endogenous SAS-6 was absent from centrioles in G1 (Figure 1C) whereas doxycycline induced SAS-6ND was expressed throughout the cell cycle (Figure 1D). In ciliated cells SAS-6ND localized to the proximal end of both mother and daughter centrioles (Figure 1E). Given the localization of SAS-6 to mother centrioles, which can function as basal bodies, we examined whether SAS-6 could affect cilia formation. Interestingly, SAS-6ND expression resulted in increased cilia length and cilia number in RPE-1 cells (Figure 1F, G and Supplementary Figure 1A). Additional experiments in human mammary epithelial cells (HMECs) and Ras-transformed MCF10AT1 showed a similar increase in ciliogenesis (Supplementary Figure 1B, C). Over-expression of WT-SAS-6 also promoted an increase in ciliogenesis, likely due to the saturation of the degradation machinery (Supplementary Figure 1B, C).

**Figure 1.**
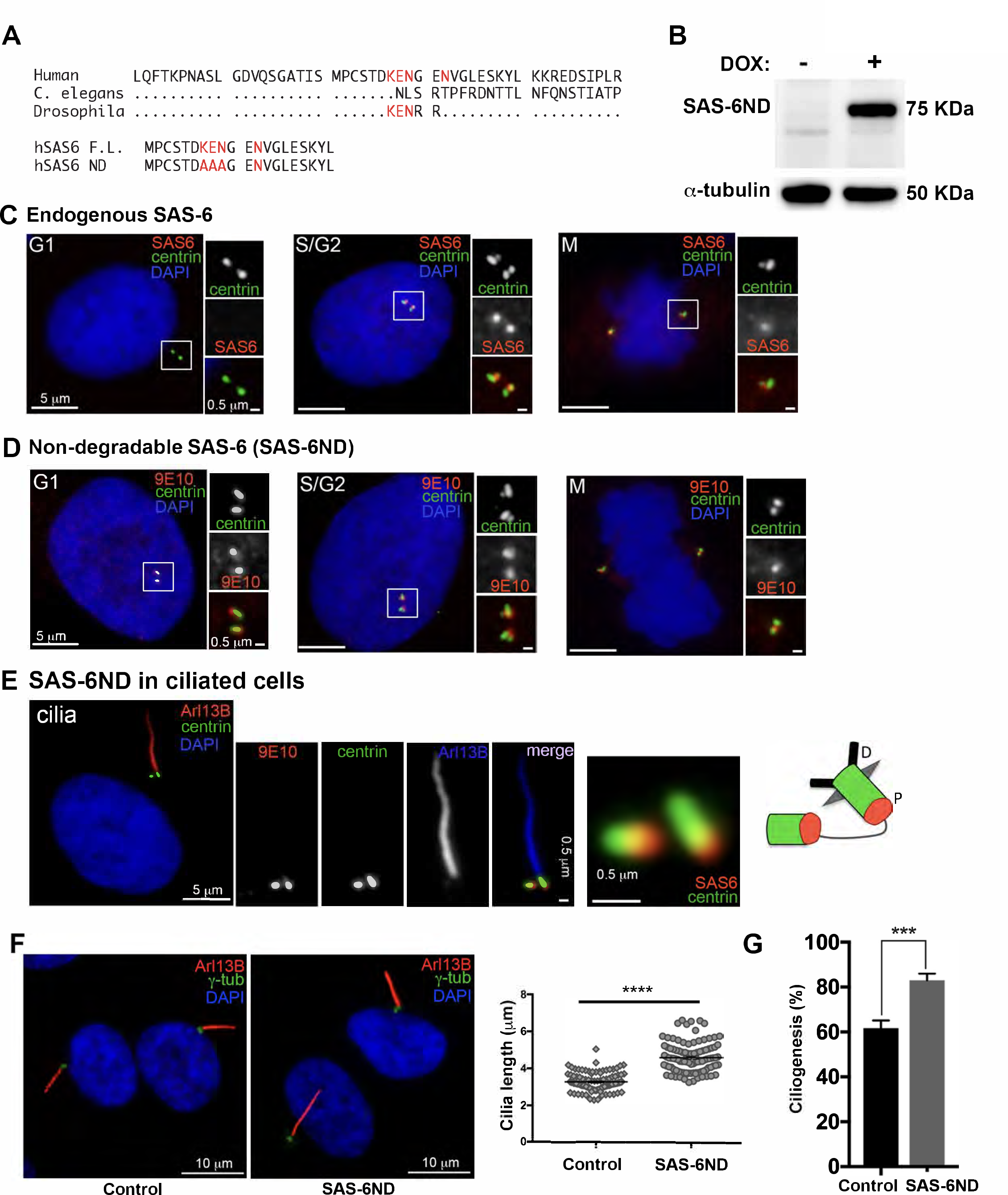
SAS-6 non degradable (SAS-6ND) is stable throughout the cell cycle and promotes increased ciliogenesis. **(A)** SAS-6 protein structure showing a stretch of human SAS-6 protein and the conserved KEN box. hSAS-6ND was generated by replacing the KEN box by an alanine stretch. **(B)** Western blot showing the expression of SAS-6ND (top) with alpha tubulin as a loading control (bottom). **(C)** Endogenous SAS-6 expression at different cell cycle stages (indicated). RPE-1 cells expressing GFP-centrin2 (green), stained with an antibody for endogenous SAS-6 (red). DNA is marked with DAPI (blue). Note that endogenous SAS-6 is absent in G1. **(D)** Non-degradable SAS-6 (SAS-6ND) expression throughout the cell cycle. Centrin2-GFP is shown in green, Myc-tagged SAS-6ND (9E10-antibody) is shown in red. DAPI is shown in blue **(E)** SAS-6ND expression in ciliated cells. The primary cilium is marked with Arl13B (red in the main panel, blue in the inset). Centrin2-GFP is shown in green, Myc-tagged SAS-6ND is shown in red. DAPI is shown in blue. An inset with both the mother centriole (basal body) and the daughter centriole is shown with a cartoon depicting SAS-6 localization (right). **(F)** Ciliation in control cells and cells expressing SAS-6ND. A quantification of cilia length is shown in the right panel. T test, p<0.0001. Note the increase in cilia length in the presence of SAS-6ND (T test, p<0.001) **(G)**. Data is representative of five independent experiments.

### SAS-6 expression is associated with poor prognosis in adrenocortical carcinoma, low grade glioma, kidney, liver and lung cancer

Centriole and cilia proteins have been shown to play a role in cancer (Bettencourt-Dias *et al*, 2011; Godinho & Pellman, 2014). Particularly, *SASS6* overexpression was found to correlate with poor prognosis in a number of tumor types (Shinmura *et al*., 2015) (Xu *et al*., 2020), highlighting the relevance of SAS-6 in cancer progression. Interestingly, knockdown of *SASS6* reduced proliferation of the highly aggressive/invasive MDA-MB-231 cell line (Du *et al*., 2021). Using recent data from The Cancer Genome Atlas (TCGA) database, we re-examined whether the expression of *SASS6* correlated with prognosis. We found that high expression of *SASS6* resulted in lower survival probability over a 10-year period in adrenocortical carcinoma, low grade glioma, renal papillary cell carcinoma and hepatocellular carcinoma (Figure 2A, B, C, D, E). Our results confirmed previous results by Shinmura et al showing that *SASS6* expression correlated with poor prognosis in kidney clear cell carcinoma and lung adenocarcinoma (Figure 2E; Supplementary Table 1). Metastatic disease is the primary cause of cancer death. Therefore, poor prognosis is often associated with increased invasion and metastasis (Ridley, 2011), suggesting that increased SAS-6 could be linked to an invasive phenotype in human cancer.

**Figure 2.**
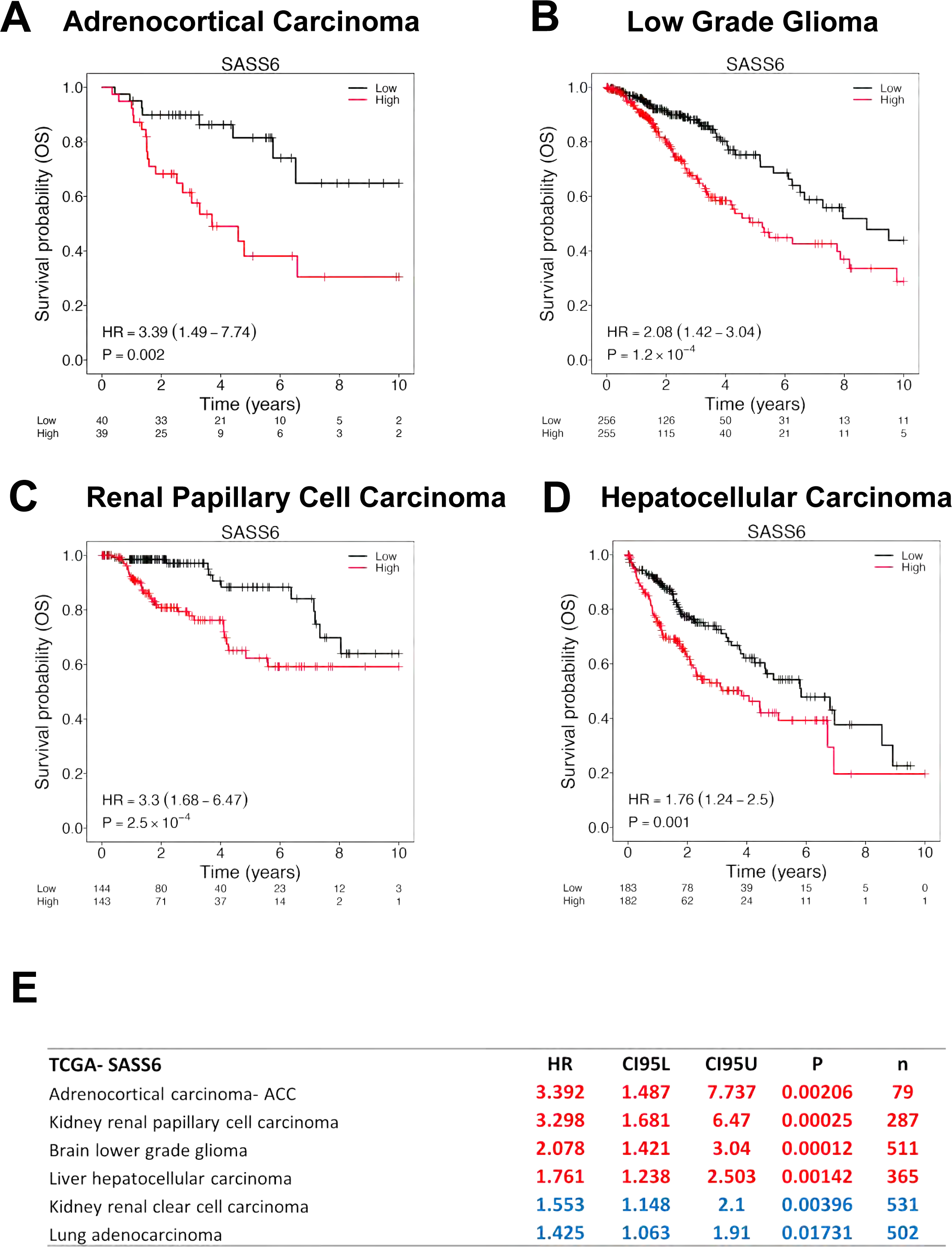
*SASS6* over expression correlates with poor prognosis. **(A, B, C, D)** TCGA analysis. Kaplan-Meier curves of patient survival probability (OS) correlated to high SAS-6 expression over time (years). High expression is indicated in red and low expression in black. The number of patients remaining in each category (low versus high) is shown below the graph. HR indicates hazard ratio, which measures the risk of having SAS-6 overexpression compared to the low expressing group. An HR higher than 1 indicates high risk. P values are shown. Shown are analyses done in TCGA adrenocortical carcinoma **(A)**, low grade glioma **(B)**, renal papillary cell carcinoma **(C)** and hepatocellular carcinoma **(D)** cohorts. **(D)** Table summarizing TCGA data analysis. CI95L, CI95U. P values and total patient numbers are shown. Note that our analysis also confirmed previous results in kidney renal clear cell carcinoma and lung adenocarcinoma.

### SAS-6 overexpression leads to invasion that depends on the presence of cilia

The ability of cancer cells to migrate to distant organs is a key process in metastasis (Yamaguchi & Condeelis, 2007) and a hallmark of cancer (Hanahan & Weinberg, 2011). Invasion requires cancer cells to leave the tissue of origin, enter blood vessels and colonize distant tissues (Yamaguchi & Condeelis, 2007). Cell migration is a multistep process initiated by cell protrusions (Ridley, 2011). Formation of protrusive structures is driven by coordinated actin polymerization at the leading edge of the cell (Yamaguchi & Condeelis, 2007). We carried out a cell protrusion assay using microporous filters (Mardakheh *et al*, 2015). Briefly, cells were seeded on top of collagen-I coated 3 μm polycarbonate transwell filters in serum-free media. After cells were attached, media in the bottom chamber was replaced with a 10% serum counterpart to signal cells to begin forming protrusions for a period of four hours. Using three different clones of cells with inducible SAS-6ND we found that upon doxycycline treatment, SAS-6ND expression showed an increase in cell protrusions in all three clones, which was calculated as a protrusion index (fold) (Figure 3A). We then carried out a three-dimensional collagen invasion assay (Gadea *et al*, 2008; Sanz-Moreno *et al*, 2008). For this experiment, cells were plated in a mixture of collagen in serum free media, spun down to the bottom of a 96-well filter and placed in an incubator at 37°C to allow collagen to polymerize. After this, media with serum was added to the top of the wells and cells were allowed to move towards serum for 24 hours before fixing them with 4%PFA. A higher number of SAS-6ND cells invaded through collagen towards serum compared to the uninduced counterpart. This was calculated as an invasion index (fold) (Figure 3B). This increase in invasion was also observed in Ras-transformed MCF10AT1 cells overexpressing both SAS-6WT and SAS-6ND (Figure 3C). Centriole and cilia signaling pathways play a role in cell migration (Kazazian *et al*, 2017; Rosario *et al*, 2015). We asked whether the invasion phenotype driven by SAS-6ND required the presence of primary cilia. To test this, we depleted SCLT1, a protein that we have previously shown to be required for ciliogenesis (Tanos *et al*, 2013) and carried out an invasion assay. Our results showed that depletion of cilia via SCLT1 removal suppressed the invasion phenotype observed upon expression of SAS-6ND (Figure 3D).

**Figure 3.**
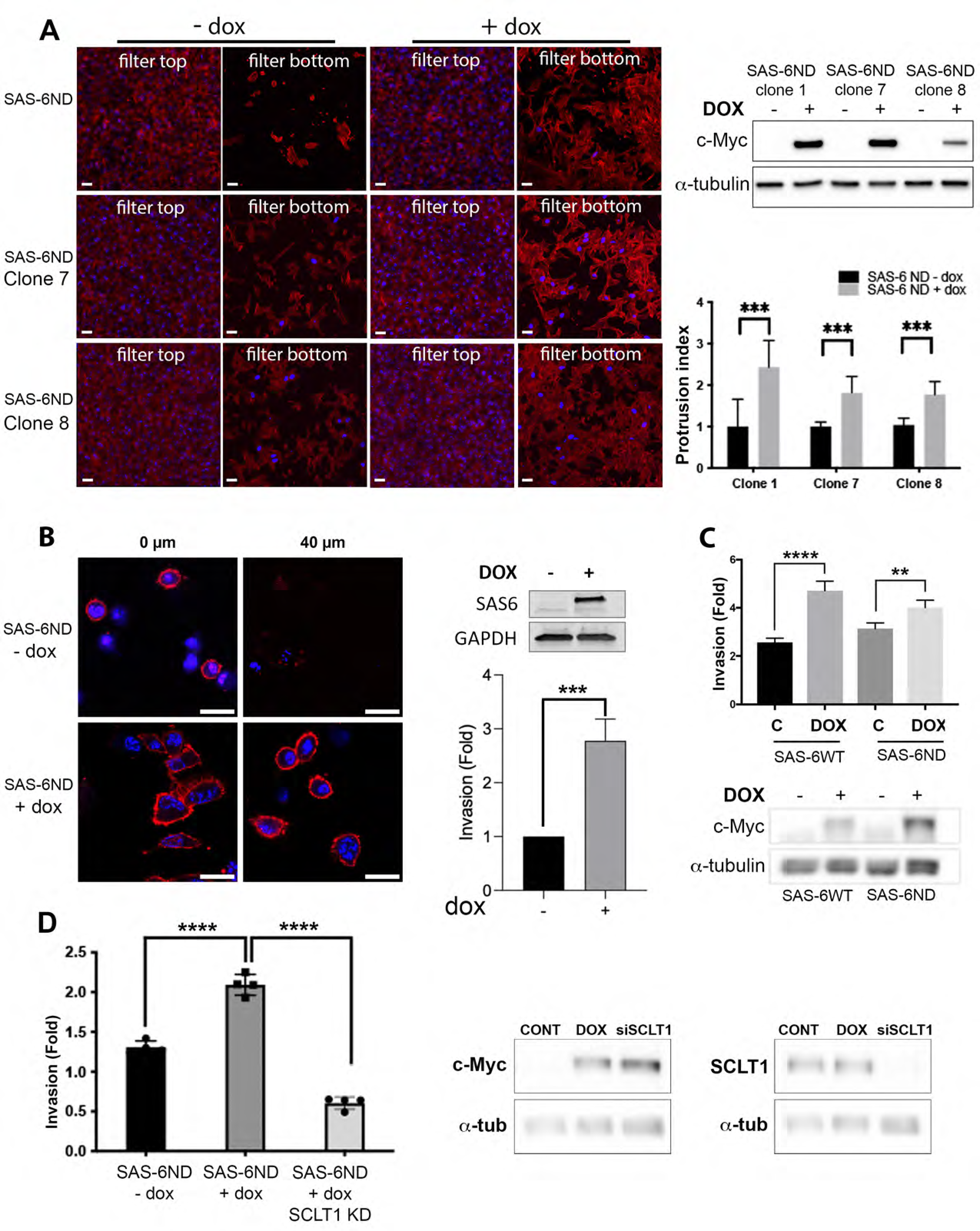
SAS-6 overexpression leads to invasion that depends on the presence of cilia. **(A)** Cell protrusion assay in cells transduced with tetracycline-inducible SAS-6ND. Confocal image of cell protrusions in 3μm transwell filters for three different SAS-6ND overexpressing clones (indicated). Panels show representative images of the top and the bottom of the filter. A control non-doxycycline treated condition is shown. The actin cytoskeleton is marked with Phalloidin (red). DAPI marks DNA (blue). Scale bar, 10μm. A western blot with the levels of Myc-tagged SAS-6ND expression is shown on the top right. α-tubulin is used as a loading control. A graph with the quantification of cell protrusions in the presence of SAS-6ND is shown. Error bars represent SD. T test is indicated, ∗∗∗p < 0.001. **(B)** Collagen invasion assay in SAS-6ND overexpressing cells. Panels show confocal images at 5 micron and 40 micron. Actin is shown in red and DNA is marked with Hoesch in blue. Scale bar, 20µm. A western blot with SAS-6ND levels is shown on the top right, GAPDH is used as a loading control. A quantification of invasion is shown in the lower right. T test, ∗∗∗p< 0.001. Data representative of 4 independent experiments **(C)** Collagen invasion assay in MCF10AT1 cells, overexpressing either SAS-6WT or SAS-6ND (indicated). A western blot with the expression of the Myc-tagged construct is shown (lower panels) with a c-Myc western blot and α-tubulin loading control (indicated). T-test, ∗∗p < 0.01, ∗∗∗∗p < 0.0001). Data representative of two independent experiments with four replicas **(D)** Collagen invasion assay in RPE-1 cells as shown in B, upon downregulation of SCLT1, a protein required to form cilia. Invasion index is shown on the right. Error bars represent SD. T test is indicated, ∗∗∗∗p< 0.001. Data representative of two independent experiments with four replicas. Western blots show the levels of SAS-6ND as well as SCLT1 levels (indicated). α-tubulin is used as a loading control.

### SAS-6 invasion-phenotype is associated with the activation of the YAP/TAZ pathway

To understand what molecular programs were activated by SAS-6 to promote invasion, we carried out transcriptomic analysis. cDNA was obtained from untreated cells and doxycycline induced cells expressing SAS-6ND and used as probes for DNA microarray hybridization. Gene set enrichment analysis (GSEA) revealed significant enrichment of genes involved in YAP/TAZ pathway activation following overexpression of SAS-6 (Figure 4A). YAP is a transcriptional coactivator that promotes TEAD-dependent gene transcription leading to increased proliferation, invasion and stem cell differentiation (Piccolo *et al*, 2023). We found that among the genes driving the enrichment of the YAP/TAZ gene set were CTGF and CYR61. Both of these genes encode matricellular proteins upregulated downstream of YAP (Piccolo *et al*., 2023) (Figure 4C). Their increased expression following induction of the non-degradable SAS-6 mutant was confirmed by qPCR (Figure 4D). This analysis showed a more than 20-fold increase in mRNA levels for cells treated with doxycycline expressing SAS-6ND (Figure 4D).

**Figure 4.**
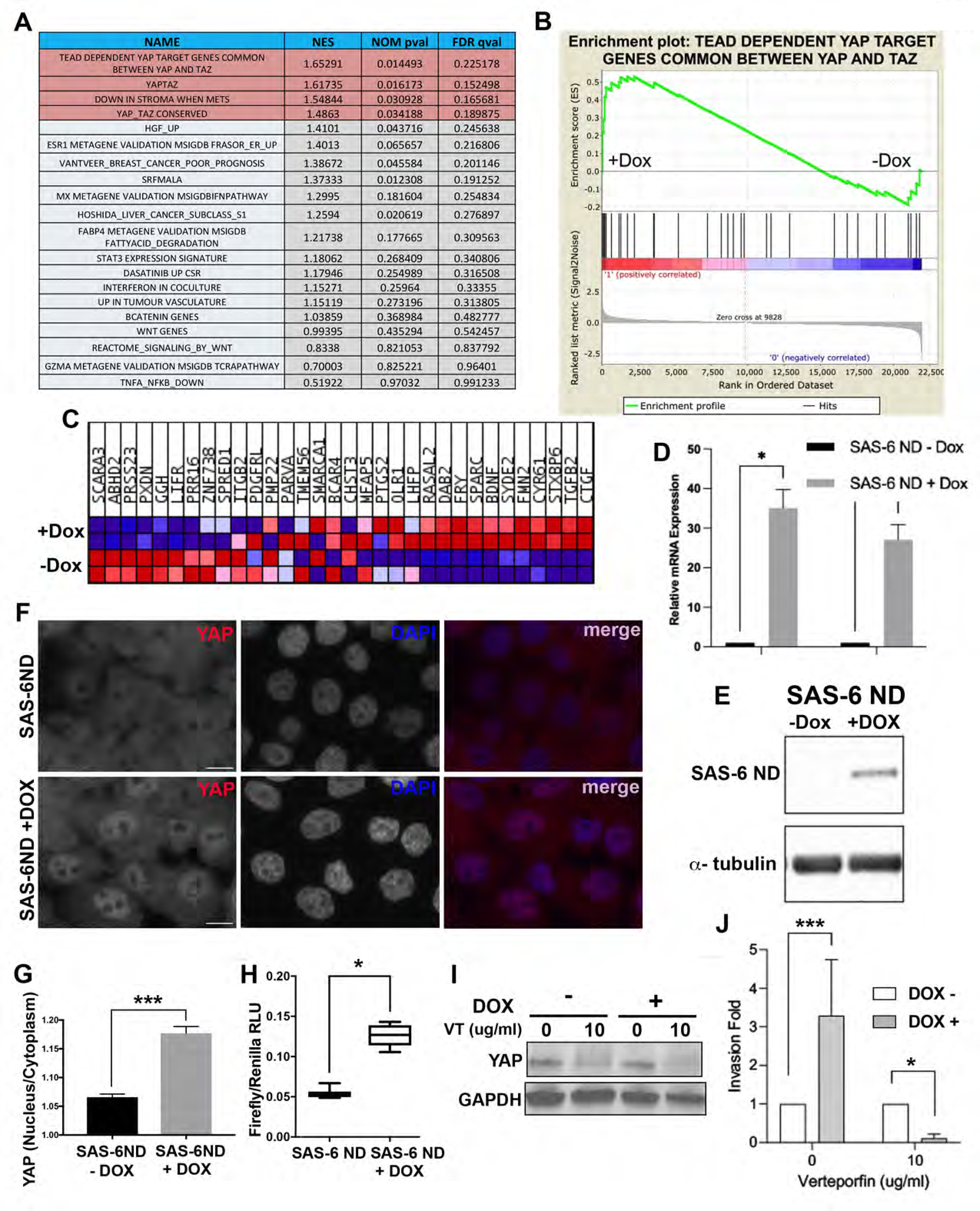
SAS-6 invasion-phenotype is associated with the activation of the YAP/TAZ pathway **(A)** Geneset enrichment analysis of microarray data. Normalized Enrichment Score (NES), Nominal P-value (NOM pval) and False Discovery Rate q-value (FDR qval) are shown for each indicated geneset. Data is representative of two independent experiments. **(B)** Enrichment plot for the dataset “TEAD DEPENDENT YAP TARGET GENES COMMON BETWEEN YAP AND TAZ” showing an enrichment score curve. NES: 1.65291. **(C)** Heat map with the list of genes driving the change. Note the increase in YAP downstream targets CTGF and Cyr61. **(D)** qPCR showing the mRNA levels of YAP/TAZ target genes, CTGF (T test, p <0.05) (left) and CYR61 (T test, p<0.05) (right). Ct values were normalized to GAPDH and expression calculated as a fold change relative to controls. Data represent mean ± SEM of three independent experiments performed in triplicate. **(E)** Western blot showing expression of SAS-6ND with and without doxycycline (indicated). **(F)** Immunostaining for YAP in cells transduced with SAS-6ND, treated with vehicle control (top) or doxycycline (lower panels). Note the increase in nuclear YAP in cells overexpressing SAS-6ND. Scale bar, 10 μm. **(G)** Quantification of the nuclear/cytoplasmic ratio in control cells (SAS-6ND - dox) and cells overexpressing SAS-6ND (+dox). T test, p <0.001. Note that SAS-6ND overexpressing cells have increased nuclear YAP. **(H)** Tuckey boxplots showing luciferase activity of YAP reporter 8xGTIIC-luc (Firefly/Renilla) indicative of YAP/TAZ activation in control or cells induced with doxycycline to express SAS-6ND (T test, p <0.05). **(I)** Western blot showing YAP protein levels upon treatment with vehicle control or verteporfin (10 μg/ml) in control cells or doxycycline-induced cells expressing SAS-6ND (indicated). **(J)** Collagen invasion assay in control or doxycycline induced cells expressing SAS-6ND upon treatment with YAP inhibitor verteporfin (10 μg/ml). N= 4 experiments

To function as a transcriptional regulator, YAP requires nuclear translocation. We estimated the extent of YAP accumulation in the nucleus by quantifying the nuclear/cytoplasmic ratio of YAP using fluorescence microscopy. SAS-6ND-expressing cells showed a marked increase of nuclear YAP (Figure 4F). Consistently, a luciferase-based reporter assay for TEAD-dependent transcription showed an increase in reporter activity upon expression of SAS-6ND (Figure 4H). Given that SAS-6 seemed to promote the activation of the YAP/TAZ pathway and considering the reported role of this pathway in cell invasion, we reasoned that blocking the YAP pathway would revert the invasion phenotype. To test this, we carried out a collagen invasion assay in the presence of verteporfin, a YAP inhibitor that disrupts YAP-TEAD interactions, decreases YAP expression and blocks transcriptional activation of downstream targets of YAP (Brodowska *et al*, 2014; Liu-Chittenden *et al*, 2012). Treatment with verteporfin decreased YAP levels in cells overexpressing SAS-6ND (Figure 4I), and completely reverted the invasion phenotype (Figure 4J), supporting that SAS-6 mediated invasion is dependent on YAP.

### Mechanism of SAS-6-regulated cell invasion

Multiple solid cancers have shown an upregulation of YAP which correlates with increased invasion, malignancy and relapse (Piccolo *et al*., 2023). Mechanical cues involving integrin activation and actin cytoskeleton contraction have been shown to increase the rate of nuclear accumulation of YAP (Piccolo *et al*., 2023). YAP accumulation in the nucleus is mediated by the opening of the nuclear pore complex (NPC) by nucleo-cytoskeletal coupling (Elosegui-Artola *et al*., 2017). However, a clear understanding of how the F-actin structure in the cytoplasm controls YAP nuclear entry remains elusive. Analysis of the actin cytoskeleton showed increased actin alignment, as measured with the orientation J plugin for image J (Figure 5A, B, C). A simple microscopic observation showed that SAS-6ND-expressing cells had a flattened morphology. Analysis of the nuclear height in these cells showed a flattened nucleus, which resulted in a decreased nuclear aspect ratio (Figure 5D, E). Further analysis showed decreased nuclear solidity and nuclear form factor and increased nuclear compactness (Figure 5F, G, H), with a (non-significant) trend to an increase in the nuclear area (Supplementary Figure 2E). Based on previous reports, nuclear deformation would support the opening of the NPC (Elosegui-Artola *et al*., 2017), which could lead to the nuclear translocation of YAP in SAS-6 overexpressing cells.

**Figure 5.**
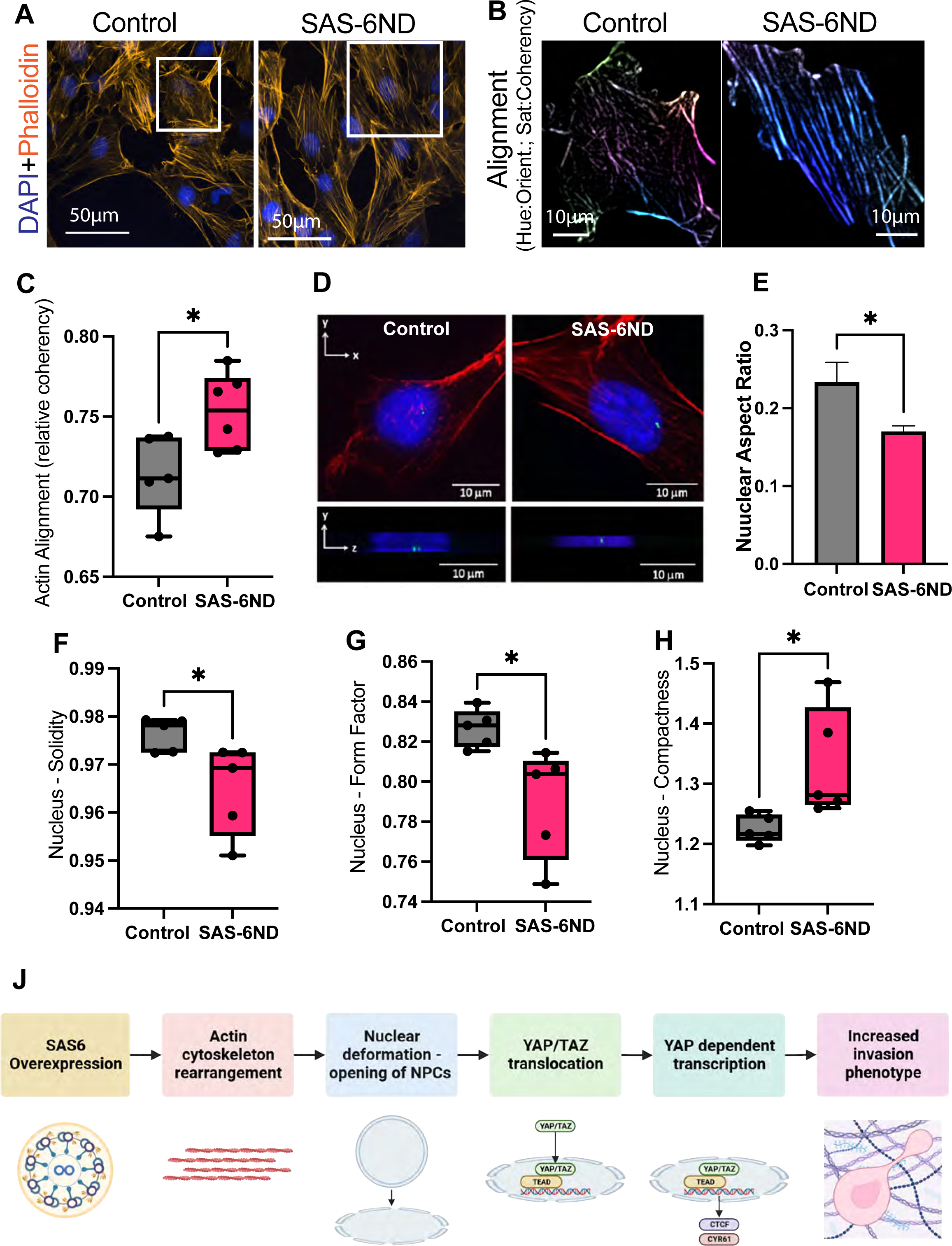
Mechanism of SAS-6-regulated cell invasion **(A)** Actin cytoskeleton staining in uninduced cells or cells expressing SAS-6ND. Phalloidin-Rhodamine is shown in orange. DNA is marked with DAPI (blue). **(B)** Actin alignment analysis showing cells from inset selection in (A). The hue is related to the orientation and the saturation is coding for the coherency. Note more aligned pixels and higher coherency values in cells overexpressing SAS-6ND (right panel). **(C)** Actin alignment quantification with Cell Profiler shows increased alignment in SAS-6ND expressing cells. N= 6 **(D)** Three-dimensional rendering of cells overexpressing SAS-6ND. Cross-section is shown where SAS-6ND expressing cells show decreased nuclear height. **(E)** Quantification of the nuclear aspect ratio for cells shown in (B), as a ratio of the vertical/horizontal axis of the nucleus (N=3). Additional Cell Profiler quantifications including nuclear solidity (as a ratio of the nuclear area/convex hull area) **(F)** and nuclear form factor (as a ratio of the area/perimeter) **(G)** show a significant decrease in SAS-6ND cells and a concomitant increase in nuclear compactness **(H)**. T test, p<0.05 for all graphs shown. N=6

Given these observations, we propose a model for SAS-6 regulation of invasion. We have found that upon overexpression of SAS-6 there are changes in the actin cytoskeleton that can lead to nuclear deformation. This nuclear deformation can lead to an increase in the nuclear import of YAP, and the activation of TEAD-dependent transcription to support cell invasion (Figure 5J).

## Discussion

Our work shows a previously unknown role of SAS-6 in the regulation of invasion. SAS-6 has been extensively characterized as a key regulator of centriole structure, supporting the initial steps of the establishment of the nine-fold symmetry and the recruitment of microtubules during centriole assembly (Pelletier *et al*., 2006) (Delattre *et al*., 2006) (Leidel *et al*., 2005) (Kitagawa *et al*., 2011; Nakazawa *et al*., 2007) (van Breugel *et al*., 2011). Recent reports suggest a potential role for SAS-6 in cancer, including a correlation between SAS-6 overexpression and poor prognosis (Shinmura *et al*., 2015) and the observation that silencing SAS-6 in breast cancer cells can suppress their proliferation (Du *et al*., 2021). SAS-6 levels oscillate throughout the cell cycle, decreasing in G1 upon targeting by the APC/Cdh1 complex via its KEN box. To maintain consistent levels of SASS6 throughout the cell cycle, we transduced cells with a SAS-6 mutant where the KEN box has been replaced with alanines (AAA), SAS-6ND (Figure 1). This SAS-6ND mutant was expressed throughout the cell cycle and displayed increased ciliogenesis and cilia length. We also observed SAS-6 localization in the basal bodies of ciliated cells (Figure 1). Interestingly, in terminally differentiated cells, such as respiratory epithelia, as well as Tetrahymena and Chlamydomonas, SAS-6 also concentrates at the base of cilia (Vladar & Stearns, 2007). This suggests that there may be other functions of SAS-6 which are yet to be uncovered. Plk4, the kinase upstream of SAS-6 has been shown to promote centrosome amplification, chromosomal instability and cancer cell invasion (Godinho & Pellman, 2014; Godinho *et al*., 2014). However, the levels of centrosome amplification with SAS-6 overexpression during our experiments are not as significant as the ones observed upon overexpression of Plk4 (Kleylein-Sohn *et al*, 2007), as SAS-6 was only expressed for 6 days.

Given the potential role of SAS-6 in cancer (Du *et al*., 2021; Shinmura *et al*., 2015), we examined the TCGA (The Cancer Genome Atlas) database in search of any clinical correlates. This analysis revealed that increased expression of *SASS6* correlated with poor prognosis in adrenocortical carcinoma, low grade glioma, renal papillary cell carcinoma and hepatocellular carcinoma (Figure 2) and confirmed previous reports of poor prognosis in kidney renal clear cell carcinoma and lung adenocarcinoma (Table 2) (Shinmura *et al*., 2015). Given that poor patient outcome has been associated with metastasis in cancer, we sought to interrogate the invasive process in cells overexpressing *SASS6*. Initial analysis in RPE-1 cells showed increased cell protrusion in collagen coated filters (Figure 1A) and increased invasion in soft agar upon expression of SAS-6ND, supporting a role for SAS-6 in invasion (Figure 3). This was confirmed in a MCF10AT1 a ras-transformed model of cancer progression by expressing either SAS-6 WT and SAS-6ND (Figure 3D). This is probably due to a saturation of the system, as the SAS-6 protein levels are higher upon doxycycline treatment even in WT-overexpressing cells (Figure 3C). Removal of cilia via knockdown of distal appendage protein SCLT1 (Tanos *et al*., 2013) reverted the SAS-6 invasion phenotype (Figure 3D), suggesting that the presence of cilia could be involved in this process. For instance, cilia-dependent signaling could promote the observed morphological changes that lead to the nuclear translocation of YAP.

It has been found that SAS6-like (SAS6L), a paralog of SAS6, localizes to the apical complex in Apicomplexa phylum, including Toxoplasma, Trypanosoma and Plasmodium (Wall *et al*, 2016). This complex is assumed to play a mechanical and secretory role during invasion in Apicomplexa to support host penetration and invasion. However, how SAS-6 contributes to invasion in this context has not been addressed.

Cell invasion is a coordinated process that requires mechanical contributions of the actin cytoskeleton as well as signaling cues (Olson & Sahai, 2009). Previous work by G. Gupta uncovered an interaction between SAS-6 and FAM21 (Gupta *et al*, 2015), a protein component of the WASH complex, which has been shown to have a direct role in Arp2/3 activation, cell protrusion, actin remodeling and invasion (Zech *et al*, 2011). Interestingly, the WASH complex assembles at centrioles (Visweshwaran *et al*, 2018), reviewed in (Fokin & Gautreau, 2021). Plk4 has been shown to regulate invasion by promoting the interaction of STIL with CEP85, STIL localization to the leading edge of the cell and Arp2/3 complex activation (Liu *et al*, 2020) (Kazazian *et al*., 2017). Given its strong interaction with STIL, this would support a role for SAS-6 in invasion. However, we only carried out SAS-6 overexpression for a maximum of 6 days, so, the level of centrosome amplification was minimal compared to the overt rosettes and *de novo* formed centrioles with amplified Plk4 (Kleylein-Sohn *et al*., 2007). Recently, Ofd1, a SAS-6 interacting protein (Go *et al*, 2021; Gupta *et al*., 2015), was shown to regulate a centriole/cilia-dependent signal for Arp2/3 complex activation, which suggests that a signal from centrioles can coordinate invasion (Cao *et al*, 2023). Future experiments will determine whether a SAS-6-mediated signal could promote direct changes in the actin cytoskeleton or an effect on the Arp2/3 complex through its interaction with the WASH complex component FAM21 (Gupta *et al*., 2015).

To understand the invasion phenotype, we examined RNA expression in SAS-6 expression in cells. Microarray and gene set enrichment analysis (GSEA) data showed that SAS-6ND promoted the activation of the YAP/TAZ pathway (Figure 4). YAP is a transcriptional coactivator promoting the transcription downstream of the TEAD promoter. The YAP/TAZ pathway has been shown to be overexpressed in cancer and promote invasion and cell proliferation (Piccolo *et al*., 2023). We validated this result using qPCR for CTGF and Cyr61 (Figure 4D), two genes downstream of YAP. Active YAP localizes to the nucleus, whereas cytoplasmic YAP is inactive and eventually phosphorylated by LATS and targeted for degradation (Piccolo *et al*., 2023). SAS-6 overexpression showed increased nuclear/cytoplasmic ratio of YAP and increased TEAD-dependent transcription using a fluorescent reporter assay in RPE-1, HMEC and MCF10AT1 cells, supporting a role of SAS-6 in the activation of this pathway. Consistently, using verteporfin, a specific YAP inhibitor, suppressed SAS-6 mediated invasion (Figure 4J). How could SAS-6 mediate this effect? To understand this phenotype we examined the morphology of the actin cytoskeleton. SAS-6ND expressing cells showed increased actin alignment and condensed stress fibers (Figure 5A, B, and C), supporting an invasive phenotype. A simple observation revealed that SAS-6 ND expressing cells appeared flattened. The YAP/TAZ pathway has been shown to be activated by mechanical force (Piccolo *et al*., 2023) and by cell flattening, which was proposed to promote the opening of the Nuclear Pore Complex (NPC) (Elosegui-Artola *et al*., 2017). Analysis of the vertical and horizontal axes in the nucleus revealed a decreased nuclear aspect ratio, thus confirming cell flattening (Figure 5D, E). Further analysis confirmed a decrease in nuclear solidity, form factor and an increase in nuclear compactness, which are consistent with the nucleus appearing flat.

Thus, we have shown that SAS-6 overexpression leads to actin cytoskeleton/morphological changes promoting a cilia-dependent invasive phenotype mediated by the YAP-pathway.

## Methods

### Cell lines and reagents

All cell lines were maintained at 37°C in a humidified atmosphere containing 5% CO2. Human telomerase-immortalized retinal pigment epithelial cells hTERT-RPE-1 or RPE-1 were purchased from American Type Culture Collection; ATCC, USA) and transduced with pBABE retro GFP-centrin 2. RPE-1 Cells were cultured in Dulbecco’s Modified Eagle Medium/Nutrient Mixture F-12 Ham (DMEM/F-12; Sigma-Aldrich, UK) media containing 0.365 mg/ml L-glutamine (Lglut), 15 mM HEPES, 1.2 mg/ml sodium bicarbonate (NaHCO3) and supplemented with 10% fetal bovine serum (FBS; Gibco, UK) and 1% penicillin/streptomycin (p/s; Gibco, UK). 1 μg/ml doxycycline was added to growth media for 6 days prior to experiments to induce SAS-6 expression. Ras-transformed MCF10AT1 cells (Miller, 1996) were kindly provided by The Barbara Ann Karmanos Cancer Institute (Detroit, MI, USA) and maintained in DMEM/F12 supplemented with 5% horse serum, 20ng/ml EGF, 0.5mg/ml Hydrocortisone, 100ng/ml Cholera toxin, 10mg/ml Insulin and 1% penicillin/streptomycin. HMECs-hTERT (Clontech) were cultured in Mammary Epithelial Cell Growth Medium with the addition of MEGM™ BulletKit (LONZA). HEK293T used for lentiviral transduction were maintained in Dulbecco’s Modified Eagle Medium (DMEM; Sigma-Aldrich, UK) supplemented with 10% FBS and 1% penicillin/streptomycin.

Stable clones of RPE-1 cells, HMECs and MCF10AT1 expressing tetracycline-inducible wild type SAS-6 or SAS-6ND were obtained via lentiviral gene transduction with the pLVX tet-on Advanced inducible gene expression system (Clontech) (Fong *et al*., 2014). Lentiviruses were produced by transfecting 293T cells with the pLVX constructs together with packaging and envelope vectors (Clontech) using the calcium phosphate precipitation method. Stable expressors were derived by selection with 5 μg/mL puromycin (Sigma-Aldrich, UK). Doxycycline was purchased from Sigma-Aldrich, UK and used at 1 μg/ml. Verteporfin (Cayman Chemical, UK) was used at 10 ug/ml.

### Western blots

Cells were lysed in ice-cold RIPA buffer (Sigma-Aldrich, UK) supplemented with a cocktail of protease and phosphatase inhibitors (Thermo Fisher Scientific, UK). Lysates were sonicated and cleared by centrifugation at 14,000 x g for 15 min. Protein content was determined using the DC Biorad Protein Assay (Biorad) following manufacturer instructions. Protein samples were subsequently denatured at 95°C for 10 min, separated by SDS-PAGE on a 4–12% polyacrylamide gradient gel and transferred to a nitrocellulose membrane. The membrane was then blocked with 5% non-fat dry milk in TRIS-buffered saline and 0.05% Tween-20 (TBST) for 1 hour before an overnight incubation at 4°C with the indicated antibodies. After this, membranes were washed 3X in TBST before and after the addition of horseradish peroxidase (HRP)-conjugated anti-mouse IgG secondary antibodies (1:2000; Cell Signalling Technology, UK; 7076) for 1 hour. Immunoreactivity was visualized using SuperSignal™ West Pico Chemiluminescent Substrate (Thermo Fisher Scientific, UK; 34080) and the BioRad ChemiDoc XRS+ imaging system.

### Immunofluorescence

Cells grown on poly-L-lysine coated coverslips were fixed in 4% paraformaldehyde (PFA) for 10 min at room temperature prior to permeabilization with 0.1% Triton X-100 in phosphate buffered saline (PBS) and blocking with 3% (w/v) bovine serum albumin and 0.1% Triton X-100 in PBS for 5 min. Primary antibodies and Alexa Fluor 594 Phalloidin (1:100; Molecular Probes, UK; A12381) were diluted to desired concentrations in blocking solution and allowed to incubate for 1 hour before three washes with 0.1% Triton X-100 in PBS (PBST). For centriolar SAS-6, PFA fixation was preceded by a 2 min permeabilization in PTEM buffer containing 20 mM PIPES (pH 6.8), 0.2% Triton X-100,10 mM EGTA and 1 mM MgCl_2_. Goat secondary antibodies conjugated to Alexa Fluor 594 (1:500 dilution; Thermo Fisher Scientific, UK) were then incubated for 1 hour followed by three PBST washes and DAPI staining (Invitrogen, UK) for DNA visualization. Coverslips were mounted with ProLong Gold antifade reagent (Invitrogen, UK).

### Antibodies

The following antibodies were used: Mouse monoclonal antibody to detect endogenous SAS-6 (1:500) was purchased from Santa Cruz Biotechnology (sc-81431). Overexpressed SAS-6 was detected with mouse anti-c-Myc (1:250; 9E10; Invitrogen, UK; 13-2500) antibody. Mouse anti-γ-tubulin (1:500; Santa Cruz Biotechnology, Inc.) and mouse anti-α-tubulin (1:2000; Sigma-Aldrich). Cilia were detected with mouse anti-acetylated-tubulin (1:2000; Sigma-Aldrich) and with rabbit anti-Arl13B (1:500; Proteintech; 17711-1-AP). Centrioles were detected with mouse anti-centrin (1:500; 3E6; Abnova; H00001070-M01). Goat secondary antibodies conjugated to Alexa Fluor 488, 594 or 680 (1:500 dilution; Thermo Fisher Scientific, UK) were used. For western blot, a Polyclonal rabbit anti-GAPDH was also used (EMD Millipore/AB2302). Western blot Secondary antibodies were horseradish peroxidase (HRP)-conjugated rabbit or mouse anti-IgG antibodies (1:2,000; Cell Signaling).

### Image acquisition, processing, and analysis

Fluorescent images were acquired on an Axio Imager M2 microscope (Zeiss, Germany) equipped with 100x, 1.4 numerical aperture (NA) oil objective; an ORCA R2 camera (Hamamatsu Photonics); and ZenPro processing software (Carl Zeiss). Images were captured with similar exposure times and assembled into figures using Photoshop (CS5, Adobe).

Deconvolution microscopy was carried out with the DeltaVision Elite (Applied Precision) using an Olympus 100x, 1.4 NA oil objective; 405 nm, 488 nm, and 593 nm laser illumination; and standard excitation and emission filter sets. Raw images were acquired using a 0.2 μm z-step size and reconstructed in three dimensions with the softWoRx 5.0.0 (Applied Precision) software. To determine the nuclear aspect ratio, height and length measurements of nuclei in the y-z plane were obtained using ImageJ.

Invasion assays were quantified in cells stained with Phalloidin and DAPI and an ImageXpress Confocal High-Content Imaging System Confocal (Molecular Devices).

Cell morphology analysis and transwell cell protrusion assays imaged on a Zeiss LSM 710 confocal microscope with a 63x NA oil objective at optimal aperture settings. 4 times averaging per image was used. Image segmentation and cell morphology analysis (as shown in Figure 5) was performed using CellProfiler, ImageJ (OrientationJ), and MATLAB as described previously (Swiatlowska *et al*, 2022). Briefly, OrientationJ produces a weighted histogram for pixels per orientation. The weight is the coherency, which is defined through the ratio of difference and sum of the tensor eigenvalues and is bounded between 0 and 1, with 1 representing highly oriented structures (Rezakhaniha *et al*, 2012).

### Ciliogenesis experiments

Cells were plated in poly-lysine-coated coverslips in 3.5-cm plates at 0.4 × 10^6^ cells per well and allowed to attach for 24 h. After this, cells were washed twice with serum-free medium, and left in serum-free medium for an additional 48 h. Cilia were detected by staining with antibodies for acetylated tubulin and ciliary membrane protein Arl13B.

### Protrusion assays and 3-D collagen invasion assays

Transwell protrusion formation assays were carried out on 3-μm pore transwell filters coated with 5 mg/ml collagen as described previously (Mardakheh *et al*., 2015). To quantify protrusions, a ratio of fluorescence intensity (measured with ImageJ) was calculated.

Collagen invasion assays were carried out as previously described (Sanz-Moreno *et al*., 2008). Briefly, cells were trypsinized and resuspended in serum-free liquid bovine type I collagen (3 mg/ml; CELLINK, UK; 5005) in DMEM, dispensed into PerkinElmer black 96-well ViewPlates and centrifuged at 1000 RPM for 5 min to force cells toward the bottom of each well. Collagen polymerization proceeded for 3 hours at 37°C, after which a final concentration of 10% FBS in DMEM/F-12 media was added to promote cell invasion into the collagen matrix. Each plate included vehicle-treated and doxycycline-treated cells as controls. Cells were fixed with a final concentration of 4% PFA for 24 h, permeabilized for 30 min with 0.5% Triton X-100 in PBS and incubated with Alexa Fluor 594 Phalloidin (1:100; Molecular Probes, UK; A12381) and DAPI for 1 hour. Confocal z-slices were captured every 10 μm (from 0-100 μm) and the number of nuclei in each plane was used to calculate an invasion index.

### Luciferase reporter assays

For the luciferase reporter assays, a TEAD-reporter construct (8xGTIIC-luciferase, Addgene, Plasmid #34615) and a CMV-Renilla (pGL4.75[hRluc/CMV], Promega) were used. Cells were seeded in 6-well plates at 70% confluency. Cells were co-transfected with 2 µg of TEAD-reporter and 100 ng of CMV-Renilla cDNA constructs using Lipofectamine 3000 according to the manufacturer’s instructions. Cells were lysed 48 hours after transfection in 100 μL of lysis buffer (Promega). Aliquots of the cell lysates were used to read luciferase emission using Dual-Glo® Luciferase Assay System (Promega) according to the manufacturer’s instructions. Reporter firefly luciferase activity was normalized to Renilla activity.

### RNA extraction

RNA was isolated from RPE-1 cells using the RNeasy mini kit (Qiagen, CA) and cDNA was generated using the High-Capacity cDNA Reverse Transcription Kit (Applied Biosystems, UK) according to the manufacturer’s protocols.

### Gene-set enrichment analyses (GSEA)

Gene expression was profiled using GeneChip™ Human Transcriptome Array 2.0. CEL file image data were converted to raw values using the R statistical language package “oligo” (Carvalho & Irizarry, 2010), available from http://www.R-project.org/. Pseudoimages were generated and inspected for artifacts. Data were normalized by Robust Multi-array Average (RMA). For Gene Set Enrichment Analysis (GSEA) (Mootha *et al*, 2003; Subramanian *et al*, 2005), the data were transformed back to linear scale from log2 values. A custom geneset matrix curated to represent cell invasion was tested for enrichment using the online GSEA module version 17 of the GenePattern platform for reproducible bioinformatics (Reich *et al*, 2006). The specific settings applied in all analyses were: Number of Permutations (1000), Permutation Type (Gene set), Enrichment statistic (Weighted), and Metric for ranking genes (t Test). Tables show the Normalized Enrichment Score (NES), nominal p-value and False Discovery Rate (FDR) q-values. The list of the specific gene sets analyzed and their sources are available in Supplementary Table 2.

### Reverse-transcription and quantitative polymerase chain reaction (qPCR)

Thermocycling was performed on the QuantStudio 7 Flex Real-Time PCR machine (Applied Biosystems, UK) using TaqMan Universal PCR Master Mix (Applied Biosystems, UK; 4366072) and predesigned TaqMan probes (Applied Biosystems, UK) for reference gene glyceraldehyde-3-phosphate dehydrogenase (GAPDH; Hs99999905) and YAP downstream targets cysteine rich angiogenic inducer 61 (CYR61; Hs00155479) and connective tissue growth factor (CTGF; Hs00170014). Relative quantification was performed according to the ΔΔCT method (relative quantification, RQ = 2-ΔΔCT) and expression levels of target genes normalized to GAPDH.

### TCGA data analysis

Pre-processed TCGA mRNA abundance and clinical data were downloaded from TCGA DCC (gdac), release: 2016_01_28. Patient groups were established by median dichotomizing SASS6 expression profile resulting in low expression and high expression groups. Cox proportional hazards model was used to estimate HR and 95% CIs with P value estimated using the log rank test.

### Statistical analysis

Statistical tests were performed with Graphpad PRISM. Statistical analyses and samples sizes are defined in the figure legends. All data are presented as mean ± standard error of mean (SEM). Paired t-tests were performed to compare two experimental conditions. P-values equal to or less than 0.05 are indicated using asterisks: *p ≤ 0.05, **p ≤ 0.01, ***p ≤ 0.001, ****p ≤0.0001.

## Acknowledgements

We thank Andrew Holland (Johns Hopkins School of medicine) for discussions about this work, Igor Vivanco (King’s College London) for critically reading this manuscript. We thank Faraz Mardakheh (Queen Mary University of London) for discussions and reagents, Khine Nyein Myint (King’s College London) and Elefterios Kostaras (Institute of Cancer Research) for reagents. BT is funded by a Kidney Research UK Senior Fellowship.

The TCGA results reported here are based, in part, upon data generated by The Cancer Genome Atlas pilot project established by the NCI and NHGRI. Information about TCGA and the investigators and institutions who constitute the TCGA research network can be found at http://cancergenome.nih.gov/.

## FIGURE LEGENDS

**Supplementary Table 1** TCGA analysis for *SASS6*.

TCGA analysis in full. Significant associations are highlighted. HR and P values are indicated.

**Supplementary Table 2** Curated genesets. Table containing geneset names, source, and the list of associated genes.

**Supplementary Figure 1.**
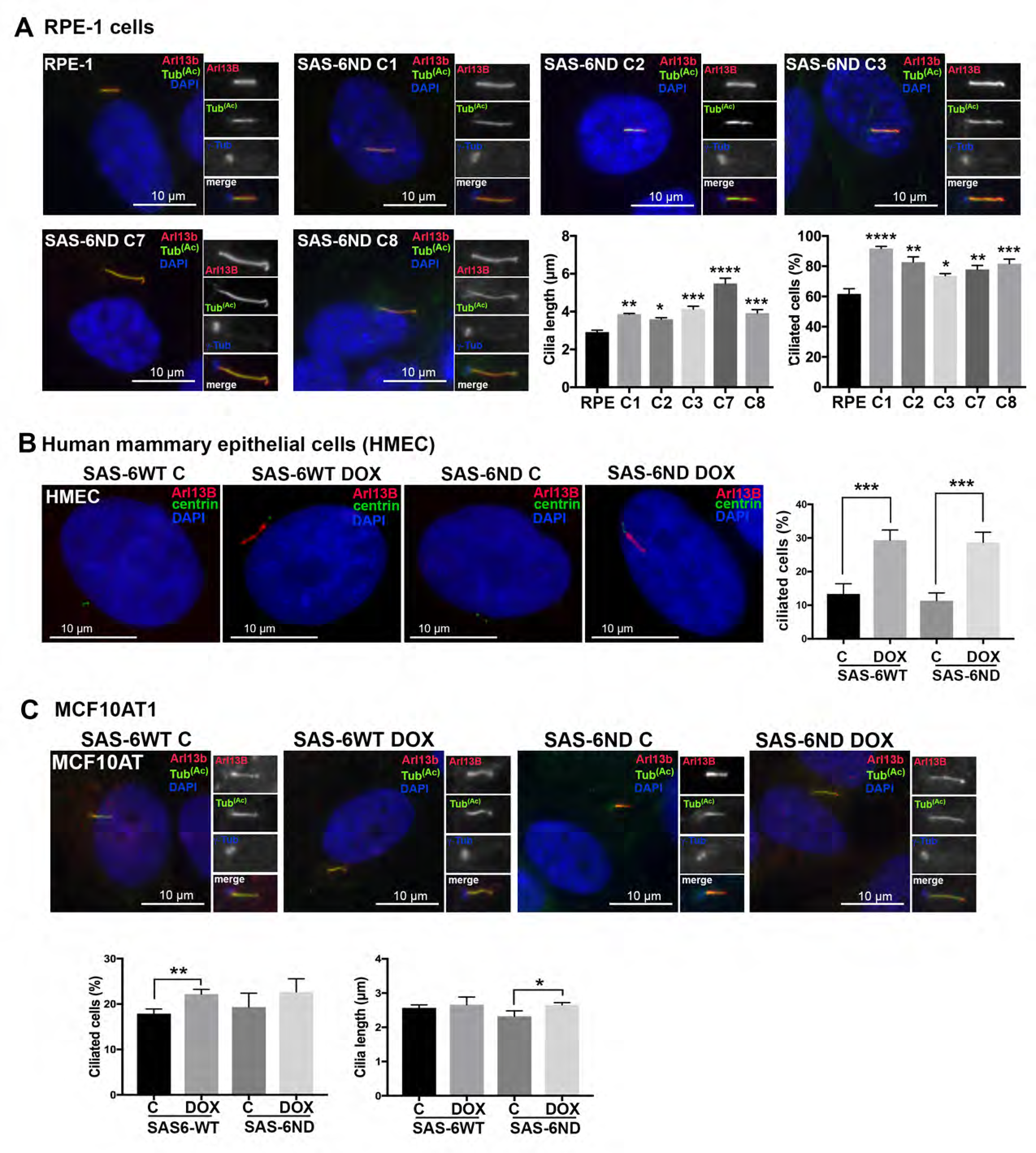
SAS-6 ND Promotes increased ciliogenesis in RPE-1 cells, HMEC and MCF10-AT. **(A)** Ciliation in control RPE-1 cells and different clones of cells overexpressing the SAS-6ND mutant (indicated as C1, C2, C3, C7 and C8). Cells were treated with doxycycline (1ug/ml) to induce expression. Cilia are marked with Arl13B (red) and acetylated tubulin (green). The centrosome is labelled with γ-tubulin (blue). DNA is marked with DAPI in blue (indicated). T-tests significance for one-way ANOVA are indicated (∗p < 0.05, ∗∗p < 0.01, ∗∗∗p < 0.001, ∗∗∗∗p < 0.0001) **(B)** Ciliation in human mammary epithelial cells overexpressing SAS-6. Cells were transduced with a stable tet-inducible vector for either wild type (WT) or non-degradable SAS-6ND. Cells were treated with vehicle (water) or doxycycline as indicated. Cilia is marked with Arl13B (red), centrin is shown in green and DNA is stained with DAPI (blue). A quantification of ciliation (percent) is shown on the right-hand side. T test, p<0.001. **(C)** Ciliation in MCF10-AT1 cells overexpressing SAS-6. Cells were transduced as in B and subjected to doxycycline treatment or control. Cilia is marked with Arl13B (red) and Acetylated tubulin (green). The centrosome is labelled with γ-tubulin (blue). DNA is marked with DAPI (indicated). A quantification of ciliated cells and cilia length is shown in the panels below. T test, p<0.01 and p<0.05 (shown).

**Supplementary Figure 2.**
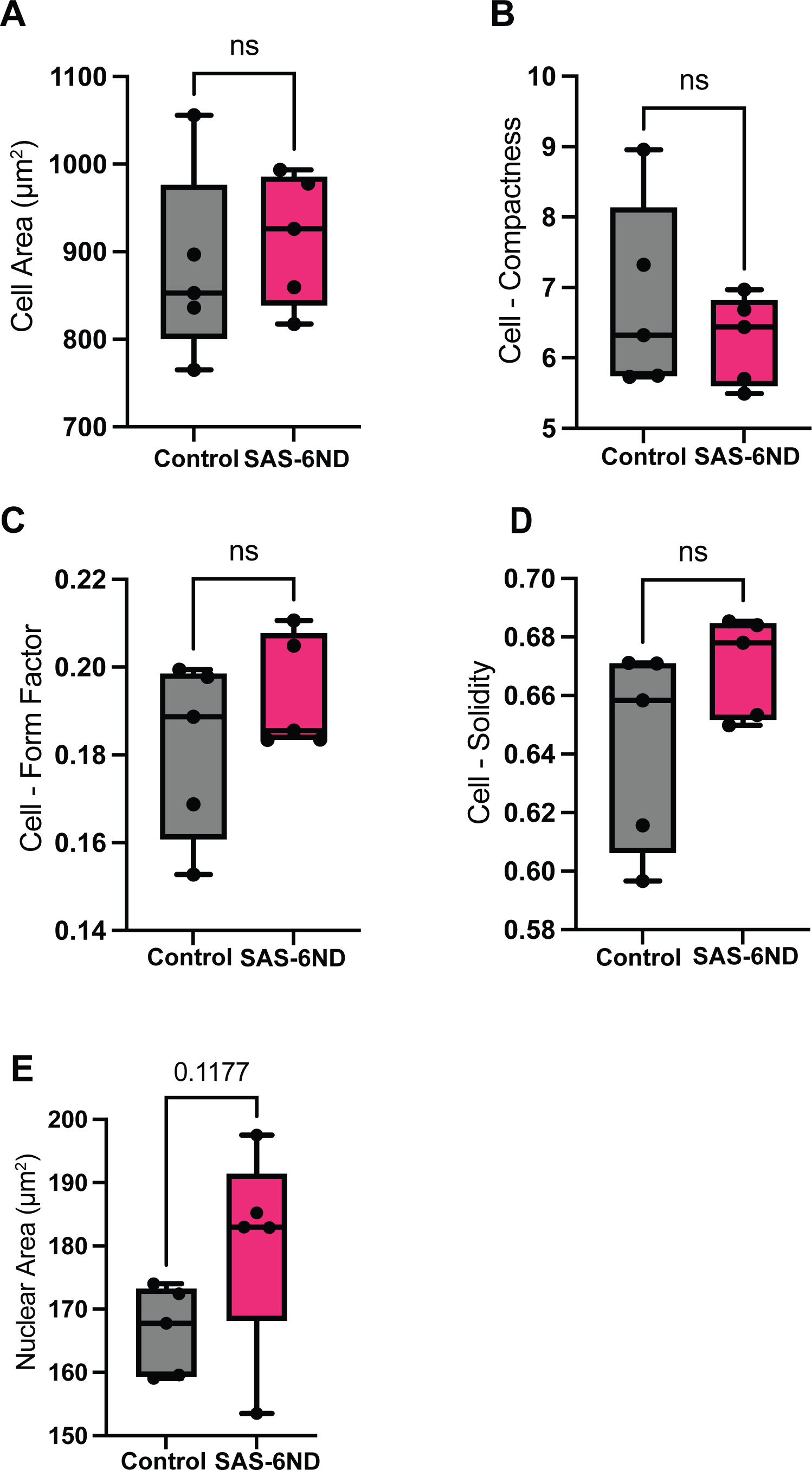
SAS-6ND overexpression does not result in overall changes in cell area. Cell profiler quantifications as described for Figure 5 in RPE-1 cells transduced with SAS-6ND, either untreated (control) or treated with 1ug/ml of doxycycline (SAS-6ND). Phalloidin staining was used to estimate additional parameters in cell profiler, including cell area (A), cell compactness (B), cell form factor (C), solidity (D) and nuclear area (E). N=6

